# A Genome-Wide Association Study Reveals a Novel Regulator of Ovule Number and Fertility in *Arabidopsis thaliana*

**DOI:** 10.1101/356600

**Authors:** Jing Yuan, Sharon A. Kessler

## Abstract

Ovules contain the female gametophytes which are fertilized during pollination to initiate seed development. Thus, the number of ovules that are produced during flower development is an important determinant of seed crop yield and plant fitness. Mutants with pleiotropic effects on development often alter the number of ovules, but specific regulators of ovule number have been difficult to identify in traditional mutant screens. We used natural variation in Arabidopsis accessions to identify new genes involved in the regulation of ovule number. The ovule numbers per flower of 189 Arabidopsis accessions were determined and found to have broad phenotypic variation that ranged from 39 ovules to 84 ovules per pistil. Genome-Wide Association tests revealed several genomic regions that are associated with ovule number. T-DNA insertion lines in candidate genes from the most significantly associated loci were screened for ovule number phenotypes. The *NEW ENHANCER of ROOT DWARFISM (NERD1)* gene was found to have pleiotropic effects on plant fertility that include regulation of ovule number and both male and female gametophyte development. Overexpression of NERD1 increased ovule number per fruit in a background-dependent manner and more than doubled the total number of flowers produced in all backgrounds tested, indicating that manipulation of NERD1 levels can be used to increase plant productivity.

**Author Summary:** Ovules are the precursors of seeds in flowering plants. Each ovule contains an egg cell and a central cell that fuse with two sperm cells during double fertilization to generate seeds containing an embryo and endosperm. The number of ovules produced during flower development determines the maximum number of seeds that can be produced by a flower. In this paper, we used natural variation in *Arabidopsis thaliana* accessions to identify regions of the genome that are associated with ovule number. Polymorphisms in the plant-specific NERD1 gene on chromosome 3 were significantly associated with ovule number. Mutant and overexpression analyses revealed that *NERD1* is a positive regulator of ovule number, lateral branching, and flower number in Arabidopsis. Manipulation of NERD1 expression levels could potentially be used to increase yield in crop plants.

## Introduction

During plant reproduction, pollen tubes deliver two sperm cells to female gametophytes contained within ovules. This allows double fertilization to occur in order to produce the embryo and endosperm in the developing seed. Angiosperms with all kinds of pollination syndromes (insect-, wind-, and self-pollinated) produce much more pollen than ovules in order to ensure successful pollination. For example, most soybean varieties produce only 2 ovules per flower, but more than 3,000 pollen grains, a 1,500-fold difference(1). Wind-pollinated plants such as maize have an even more extreme difference in pollen production vs. ovule production per plant, with more than 1 million pollen grains versus an average of 250 ovules per plant (a 4000-fold difference(2)). *Arabidopsis thaliana*, which is a self-pollinating plant, also produces an excess of pollen, with at least 2000 pollen grains per flower compared to an average of 60 ovules per flower(3). Since pollen is produced in excess, in self-pollinated plants the number of ovules (i.e. female gametes) sets the maximum seed number per flower.

The ability to manipulate ovule number to increase the reproductive potential of plants requires an understanding of the molecular pathways that control ovule initiation. The model plant *Arabidopsis thaliana* produces flowers with four whorls of organs: sepals, petals, stamens, and carpels. The inner whorls (3 and 4) are responsible for sexual reproduction, with pollen (the male gametophytes) produced in the whorl 3 stamens and the female gametophytes (also known as the embryo sacs), produced in ovules contained within the whorl 4 carpels. Specification of the 4 whorls is controlled by the “ABC” genes, with the C-class gene AGAMOUS (AG) a major regulator of carpel development(4).

In Arabidopsis, ovules are initiated from the carpel margin meristem (CMM) at stage 9 of floral development(5). The Arabidopsis gynoecium comprises two carpels that are fused vertically at their margins (6). The CMM develops on the adaxial face of the carpels (inside the fused carpel cylinder) and will give rise to the placenta, ovules, septum, transmitting tract, style, and stigma. Once the placenta is specified, all of the ovule primordia are initiated at the same time(7). Subsequently, each primordium will be patterned into three different regions: the funiculus, which connects the ovule to the septum; the chalaza, which gives rise to the integuments; and the nucellus, which gives rise to the embryo sac. Ovule development concludes with the specification of the megaspore mother cell within the nucellus which undergoes meiosis followed by three rounds of mitosis to form the mature haploid embryo sac (reviewed in(8)).

In Arabidopsis, CMM development requires coordination of transcriptional regulators involved in meristem function with hormone signaling (reviewed in(6)). Most mutants that have been reported to affect ovule number have pleiotropic effects related to the establishment of polarity and boundaries during gynoecial development (reviewed in (9)). For example, the *AINTEGUMENTA* (*ANT*) transcription factor regulates organ initiation and cell divisions during flower development(10). *ANT* acts redundantly with the related gene *AINTEGUMENTA-LIKE6/PLETHORA3* to regulate carpel margin development and fusion which leads to a modest reduction in ovule number. This phenotype is exacerbated when *ant* is combined with mutations in other carpel development transcriptional regulators, such as *SEUSS (SEU), LEUNIG (LUG), SHATTERPROOF1* and 2 (*SHP1* and *SHP2*), *CRABSCLAW*(CRC), *FILAMENTOUS FLOWER* (*FIL*), and *YABBY3* (*YAB3*). Mutant combinations between *ant* and these mutants leads to severe defects in carpel fusion coupled with severe reductions in the marginal tissues that give rise to the CMM (summarized in(6)). An extreme example is the double mutant *seu-3 ant-1* which results in a complete loss of ovule initiation due to defects in CMM development(11). The organ boundary genes, *CUP-SHAPED COTYLEDON1* and *2* (*CUC1* and *CUC2*), are also required for CMM development and subsequent ovule initiation. *ant cuc2* mutants with *cuc1* levels decreased specifically in the CMM by an RNAi construct driven by the *SEEDSTICK* promoter show an 80% reduction in ovule number, indicating that *ANT* controls cell proliferation while *CUC1/2* are necessary to set up the boundaries that allow ovule primordia to be initiated(12).

Plant hormones are also involved in gynoecium development and can have both indirect and direct effects on ovule number. Auxin biosynthesis, transport, signaling, and transport mutants have varying effects on gynoecium development and patterning, many of which lead to pleiotropic effects on tissues and organs derived from the CMM(13). Treatment of developing flowers with the auxin polar transport inhibitor NPA showed that an apical-basal auxin gradient in the developing gynoecium is necessary for patterning events that lead to ovule initiation(13). Cytokinin has also been implicated in ovule initiation and development in Arabidopsis. Notably, triple mutants in the *ARABIDOPSIS HISTIDINE KINASE* (*AHK*) cytokinin receptors, *AHK2, AHK3*, and *AHK4/CRE1*, displayed a 90% reduction in ovule number due to decreased cytokinin signaling(14). Conversely, double mutants in the cytokinin degrading cytokinin oxidases/dehydrogenases (CKXs) displayed higher cytokinin levels in inflorescences and produced more than double the number of ovules as in wild-type controls(15). Brassinosteroids (BR) may also be positive regulators of ovule initiation. Gain-of-function mutants in the BR-induced transcription factor *BZR1* had increased ovule number per flower while BR-deficient and insensitive mutants had decreased ovule number compared to wild-type controls. Upregulation of the ovule regulators *ANT* and *HULLENLOS* (*HUL*) was correlated with *BZR1* activity, indicating the BR signaling positively regulates ovule development(16). Recently, gibberellins were shown to negatively regulate ovule number independently of auxin transport and signaling. A gain-of-function mutation in the DELLA gene, GA-INSENSITVE (GAI) and loss-of-function mutations in GA receptors all led to increased ovule number in Arabidopsis (17).

To date, research on the factors controlling ovule number has been dominated by the analyses of mutants that have been identified based on pleiotropic effects on gynoecium development. In an attempt to identify loci that regulate ovule number without affecting other aspects of flower morphology, we took advantage of natural variation in ovule number in Arabidopsis accessions from diverse geographical locations. Over 7,000 natural accessions are now available with intraspecific variation, and next generation sequencing has been used to generate data on single nucleotide polymorphisms (SNPs) from over 1,000 of these accessions as part of the 1001 genomes project(18). 100s of different phenotypes have been analyzed in this collection such as flowering time, leaf shape and size, the ability to resist to pathogens, etc.(19).

In this study, we identified variation in ovule number per flower in a screen of 189 Arabidopsis accessions and conducted a Genome Wide Association Study (GWAS) to identify loci associated with the ovule number trait. Further analysis of two loci identified in our GWAS revealed that the *NEW ENHANCER of ROOT DWARFISM (NERD1)* and *OVULE NUMBER ASSOCIATED 2 (ONA2)* genes participate in the determination of ovule number during Arabidopsis flower development. The discovery of new ovule number regulators in Arabidopsis has the potential to provide targets for the development of crop varieties with greater yield.

## Results

### Natural Variation in Ovule Number

We set out to identify new regulators of ovule number in Arabidopsis by taking advantage of phenotypic variation in naturally occurring accessions. We obtained 189 Arabidopsis accessions from the ABRC and assayed them for variation in ovule number (Table S1). Since ovule number can vary throughout the life cycle of the plant(20), we determined the average number of ovules from flowers 6-10 on the main stem of plants that were vernalized for 4 weeks and then grown in long days at 22°C. Under these growth conditions, accessions displayed a remarkable diversity in ovule number per flower, with a range of 39-82 ovules per flower (Fig 1A-B). The commonly used reference accession, Col-0, falls in the middle of the range with an average ovule number of 63±3 ovules.

**Fig 1.**
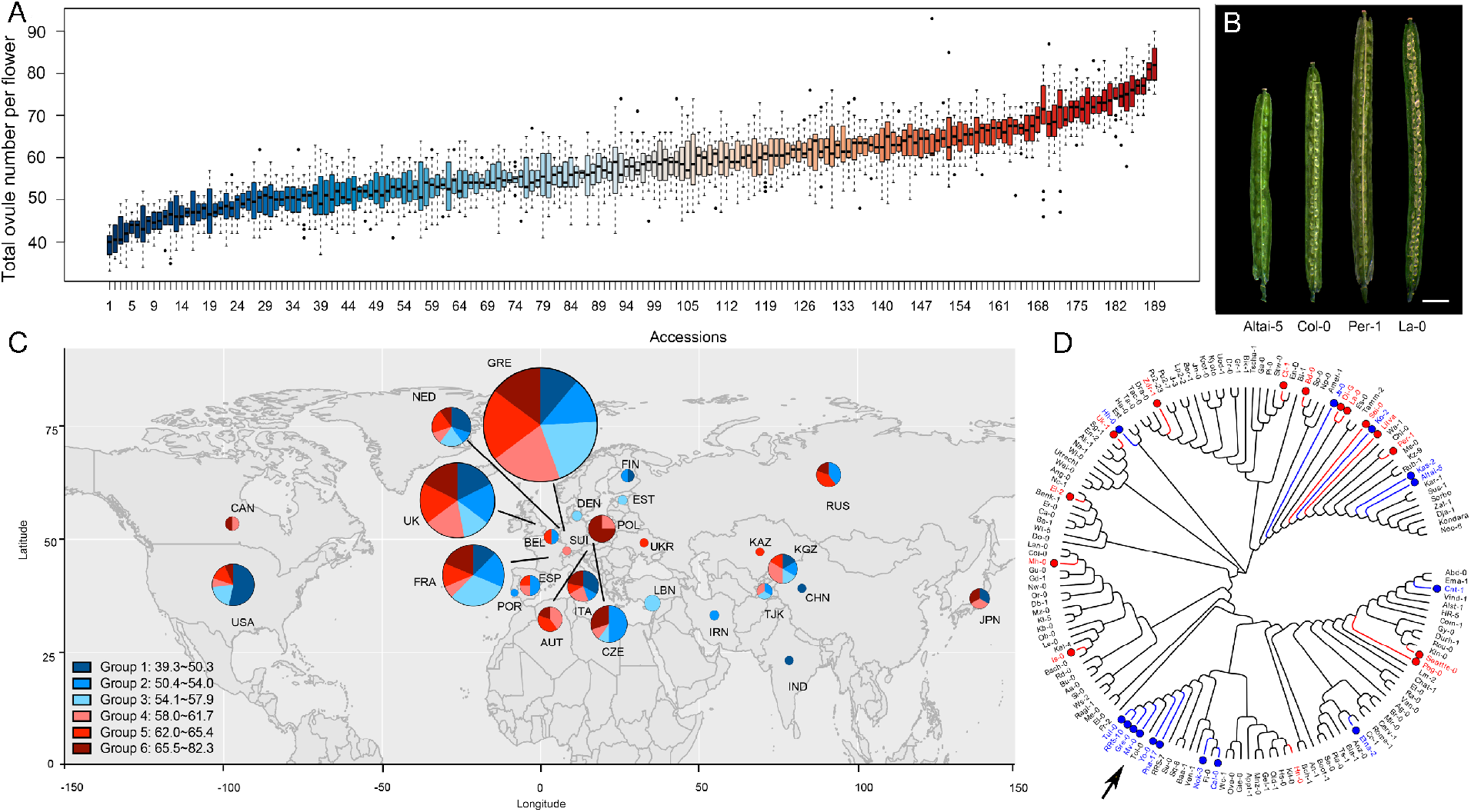
Arabidopsis accessions display natural variation in ovule number per flower. (A) Boxplot of ovule number in 189 Arabidopsis accessions. The detailed names of these accessions are in Supplementary Table 1. (B) Dissected siliques of low, medium, and high ovule number accessions (Bar=2.0mm). (C) Geographic distribution of accessions. Groups 1 to 6 correspond to ovule number ranking (from low to high), with the colors corresponding to panel A. The pie chart indicates the percentage of accessions in each country in each ovule number group, while the size of the pie chart corresponds to the total number of accessions per country. (D) Cladogram of accessions used in this study (generated in MEGA7). The 15 accessions labeled in blue had the lowest ovule numbers, and the 15 accessions labeled with red had the highest ovule numbers. The arrow points to a clade with clustered low ovule number accessions.

In contrast to flowering time variation which has been shown to correlate with latitude of origin in Arabidopsis accessions(21), ovule number was not strongly correlated with location of origin in the accessions analyzed (Fig 1C, S1). Mapping ovule number data onto a cladogram of the accessions used in our study revealed a cluster of low ovule number accessions in one specific clade, indicating that these closely-related accessions may have similar genetic control of ovule number (Fig 1D). Interestingly, several clades were made up of accessions with high, medium, and low ovule numbers. This suggests that the ovule number trait may be regulated by different loci that have been selected for in some lineages.

### GWAS reveals SNPs linked to natural variation in ovule number

In order to identify genomic regions linked to variation in ovule number, we assessed whether the average ovule number per flower from 148 accessions was predicted by Single Nucleotide Polymorphisms (SNP) available in the 1001 Full-sequence dataset; TAIR 9. Logarithmic transformation was applied to the ovule number data to make the results more reliable for parametric tests. Associations were tested for each SNP using a linear regression model (LM) and the results were analyzed using GWAPP(22) (Fig 2A). A significance cutoff value of −log_10_(*p* values) ≥ 6.2 identified at least 9 genomic regions that are associated with variation in ovule number, while a higher cutoff of −log_10_(*p* values) ≥ 7.5 identifies only four significant genomic regions.

**Fig 2.**
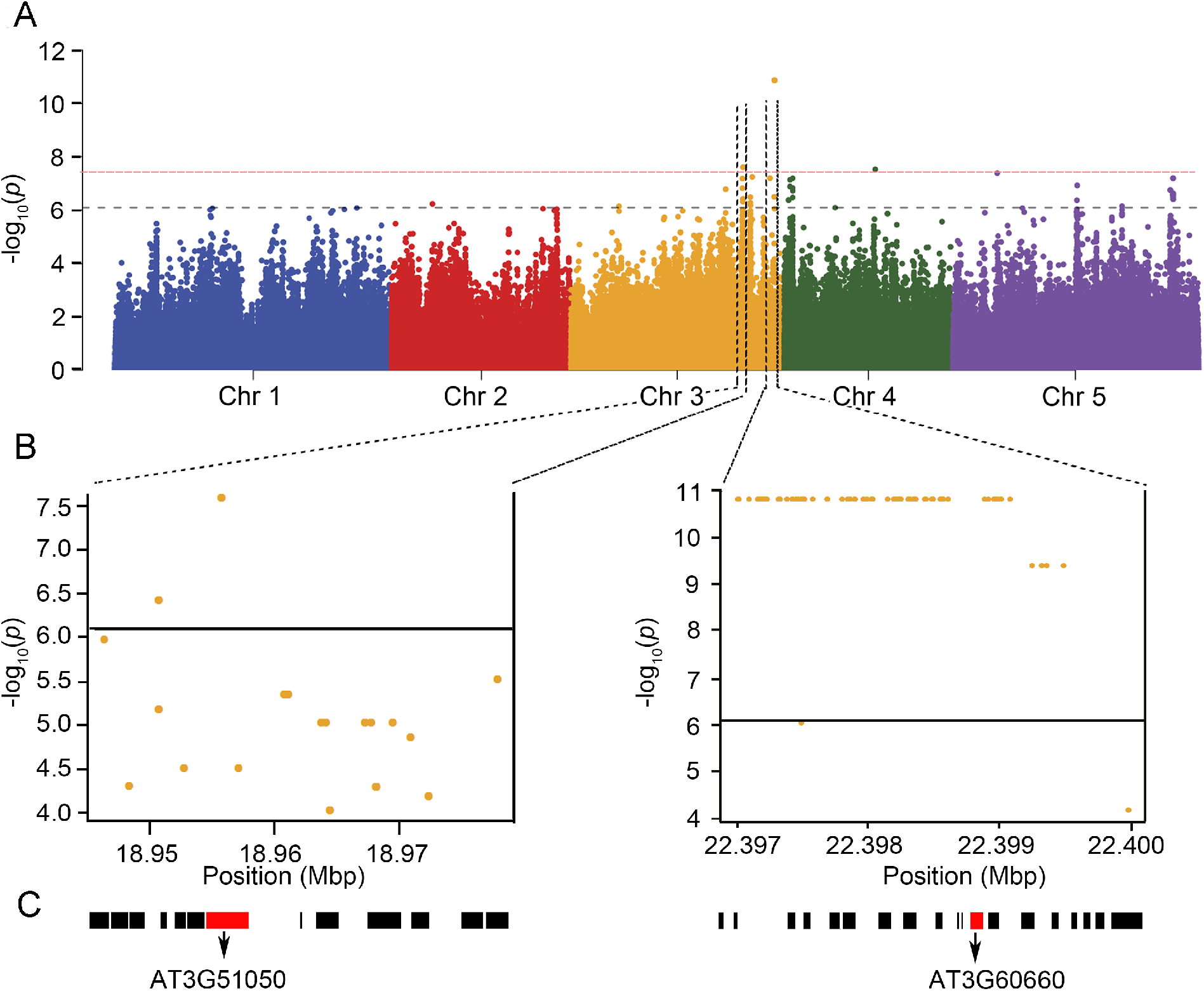
GWAS identifies candidate loci associated with ovule number per flower. (A) Manhattan plot for the SNPs associated with ovule number per flower. Chromosomes are depicted in different colors. The horizontal blue dashed line corresponds to a −log_10_(p values) ≥ 6.2 and the red dashed line corresponds to −log_10_(*p* values) ≥ 7.5 after Benjamini Hochberg (False Discovery Rate) correction. (B) The genomic region surrounding the two most significant GWA peaks on Chromosome 3. (C) Genes in the genomic region surrounding the two significant GWA peaks in (D). The red boxes are AT3G51050 (*NERD1*) and AT3T60660 (*ONA2*).

We next determined if known ovule number regulators (9, 15, 17, 23) colocalized with our GWAS loci. Of the previously described loci, only BIN2 mapped close to a significant association (Fig S2). This indicates that our GWAS has identified novel functions for loci in the regulation of ovule number. For further analysis, we focused on the two loci with the lowest p-values which are both located on the long arm of chromosome 3 (Fig 2B-C). Genes containing the most significantly associated SNPs as well as the 10 surrounding genes around the highest peak were considered as candidates for regulating ovule number. These genes were further prioritized based on whether they are expressed in developing pistils, by examining publicly available transcriptome data in ePlant(24) (Fig S3). Based on this prioritization, 35 candidate genes were selected, and of these 26 insertion mutants were available from the Arabidopsis Biological Resource Center. All 26 mutants were evaluated for changes in ovule number compared to the wild-type background, Col-0 (Table S2). Of these, two had significantly reduced ovule number compared to the Col-0 control (Fig 3A, S4). The strongest ovule number phenotype was found in insertion mutants in At3g51050, a gene that was recently identified in a screen for enhancers of exocyst-mediated root phenotypes and named *NEW ENHANCER OF ROOT DWARFISM 1* (*NERD1*)(25). The second locus with T-DNA insertions affecting ovule number identified in our screen was At3g60660, which we call *OVULE NUMBER ASSOCIATED 2* (*ONA2*). *ONA2* encodes an unknown protein containing a DUF1395 domain (TAIR) (Fig 2C).

**Fig 3.**
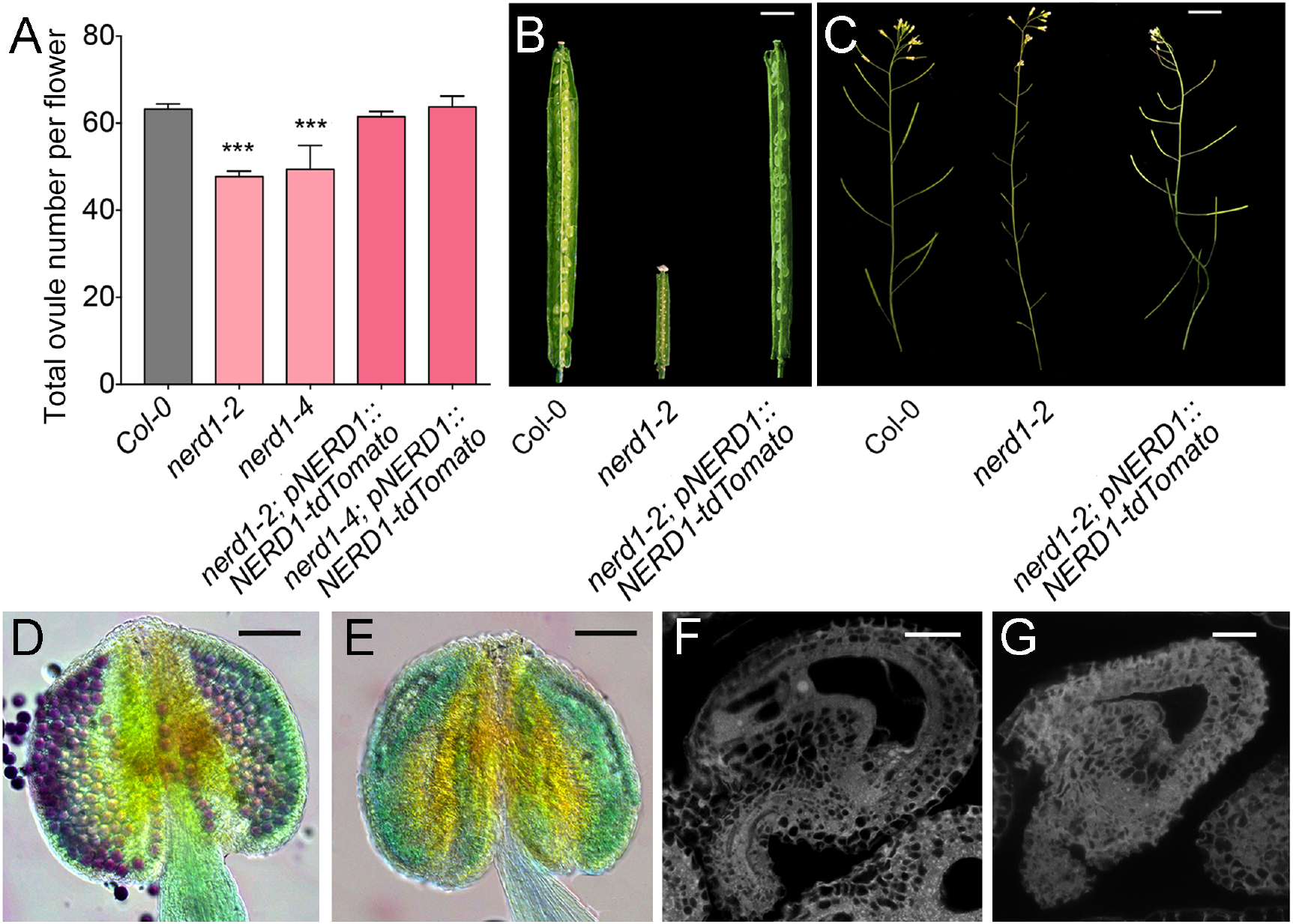
*nerdl* mutants have reduced fertility. (A) Bar graph of ovule number per flower in Col-0, *nerd1-2, nerd1-4* and their complementation lines. “***” indicates statistical significance (*p* value<0.001 determined by Student’s t-test). (B-C) *nerd1-2/nerd1-2* displays lower ovule number, infertile ovules, and short siliques compared to Col-0 and complemented lines. (D) Alexander stained Col-0 anthers have viable pollen grains (red indicates viable pollen while the green is non-viable). (E) Alexander-stained *nerd1-2/nerd1-2* anthers have no viable pollen. (F) Col-0 mature ovule with a normal embryo sac. (G) *nerd1-2/nerd1-2* mature ovule with an aborted embryo sac. Bars=2.5mm (B-C), 20μm (D-G).

### *NERD1* is a positive regulator of ovule number

We focused our analysis on *NERD1* since it had the strongest effect on ovule number. The fruits of homozygous mutants in two T-DNA insertion alleles, *nerd1-2* (described in (25) and *nerd1-4*, had fewer ovules than wild-type plants (Fig 3A). To confirm that this resulted from the disruption of NERD1, we generated transgenic plants expressing a translational fusion of the *NERD1* protein with the red-fluorescent protein tdTomato under the control of the native *NERD1* promoter. In *nerd1-2* and *nerd1-4* mutants, this transgene fully rescued the reduced ovule phenotype and resulted in plants with ovule numbers indistinguishable from Col-0 (Fig 3A). This demonstrates that *NERD1* is a positive regulator of ovule number.

In addition to reduced ovule number per flower, homozygous *nerd1* mutants had fertility defects. Homozygous *nerd1-2* mutants had 100% unfertilized ovules and homozygotes of the less severe *nerd1-4* allele had 75% unfertilized ovules (Fig 3B-C and S5). *nerd1-2* mutants displayed both male and female defects. No viable pollen grains were produced (Fig 3D-E) and embryo sacs were often aborted, leading to 40% unfertilized ovules in *nerd1-2/nerd1-2* plants pollinated with wild-type pollen (Fig 3F-G and S5). We performed reciprocal crosses between heterozygous *nerd1-2* mutants and Col-0 to determine if the fertility defects were gametophytic or sporophytic. When heterozygous *nerd1-2/NERD1* was used as the female, there was no transmission defect, demonstrating that the reduced female fertility in *nerd1-2* mutants was not gametophytic. When *nerd1-2/NERD1* was used as the pollen donor, the transmission efficiency of the mutant allele was reduced to 45%, indicating a male gametophytic transmission defect (Table 1).

**Table 1.** Transmission efficiency of the *nerdl-2* allele determined by reciprocal crosses with wild-type Col-0.

| Female | Male | *NERD1*/<br>*NERD1* | *nerd1-2*/<br>*NERD1* | Female TE% | Male TE % |
| --- | --- | --- | --- | --- | --- |
| *nerd1-2/NERD1* | Col-0 | 43 | 42 | 97.67 | NA |
| Col-0 | *nerd1-2/NERD1* | 92 | 41 | NA | 44.57 |

### *NERD1* encodes an integral membrane protein

Phylogenetic analyses indicate that *NERD1* is a member of a low-copy number, highly conserved gene family that is found throughout the plant kingdom and in cyanobacteria (Fig 4A-B and (25)). The NERD1 protein is predicted to be an integral membrane protein with a signal peptide and one transmembrane domain (Fig 5A). The majority of the protein is predicted to be extracellular, with the transmembrane domain located near the C-terminus and a 17 amino acid cytoplasmic extension. Transient expression of a NERD1-GFP fusion in *Nicotiana benthamiana* with subcellular markers confirmed that NERD1 co-localizes with the Golgi marker and partially colocalizes with a plasma membrane marker (Fig 5B and D). NERD1 does not co-localize with ER, peroxisome, and plastid markers (Fig 5C and S6).

**Fig 4.**
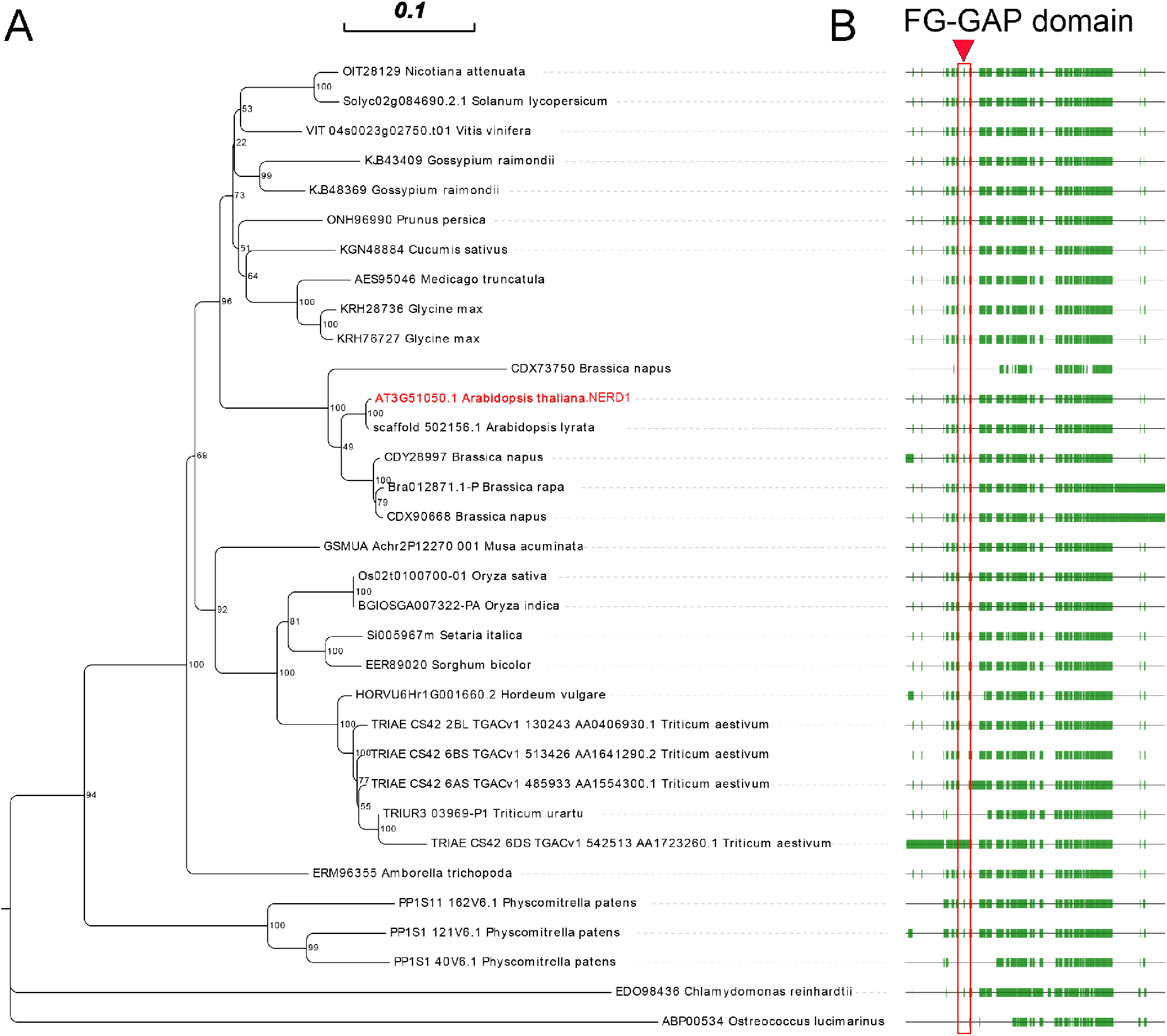
NERD1 is conserved across plant lineages. (A) Phylogenetic tree from amino acid alignment of NERD1 generated using the Neighbor-joining method in MEGA7. (B) NERD1 alignment showing sequence conservation among species. The green boxes correspond to conserved regions of NERD1 and the lines represent gaps in the alignment. The red box indicates the FG-GAP domain annotated in NERD1 by Langhans, et al., 2017.

**Fig 5.**
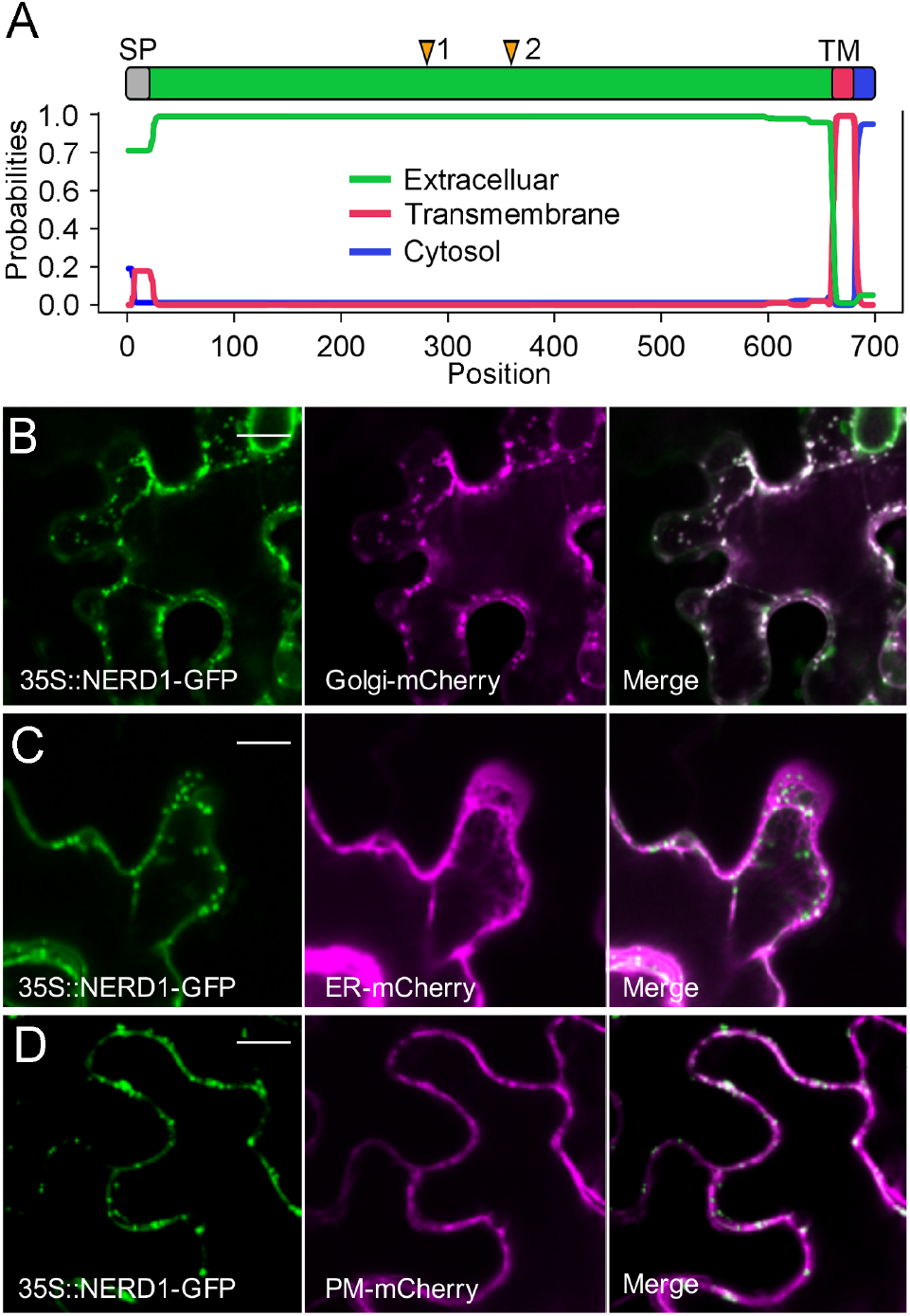
NERD1 co-localizes with a Golgi marker in *N. benthamiana* epidermal cells. (A) NERD1 protein domains determined by TMHMM 2.0. (B) NERD1-GFP (green signal) co-localizes with Golgi-mCherry (magenta signal) in *N. benthamiana* epidermal cells. (C) NERD1-GFP (green signal) does not co-localize with ER-mCherry (magenta signal). (D) NERD1-GFP (green signal) partially co-localizes with PM-mCherry (magenta signal). Bars=15 μm.

### *NERD1* is expressed throughout Arabidopsis development

The *nerd1* ovule number and fertility phenotypes suggest that *NERD1* should be expressed in developing flowers. We used a *NERD1_pro_::gNERD1-GUS* fusion to examine *NERD1* expression throughout Arabidopsis development. In *NERD1_pro_::gNERD1-GUS* inflorescences, GUS activity was detected throughout flower development, including inflorescences, developing and mature anthers, and in the stigma, ovules, and carpel walls of mature pistils (Fig 6). NERD1-GUS activity was present in the carpel margin meristem (CMM) in stage 9 flowers, where ovule initiation occurs (Fig 6C and E). *NERD1* reporter expression in the CMM during pistil development is consistent with a role for *NERD1* in ovule initiation. During seedling development, the *NERD1-GUS* reporter was detected in shoot and root apical meristems (SAM and RAM) and in the vasculature (Fig 6I).

**Fig 6.**
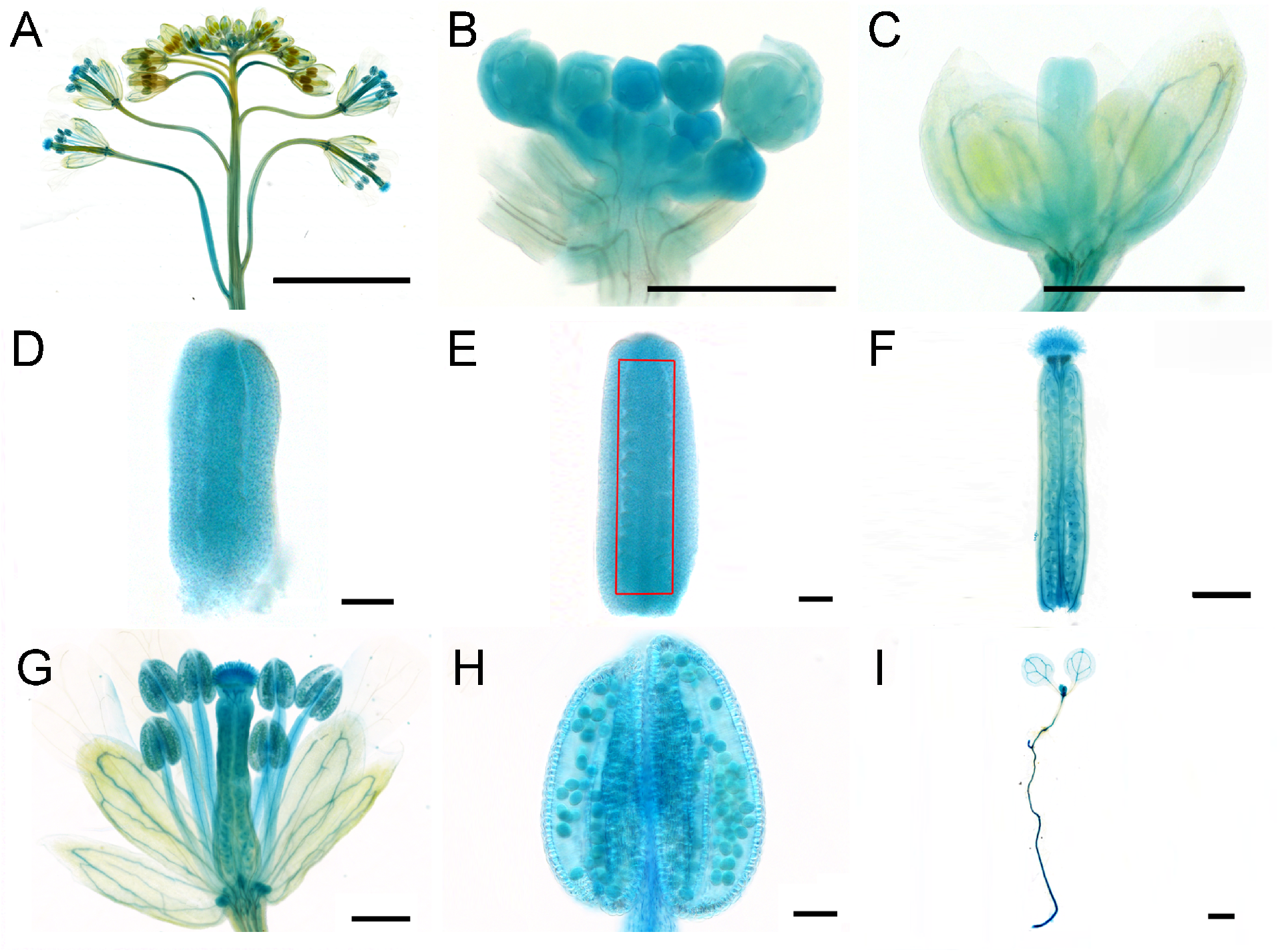
NERD1 expression in plant development. The NERD1_pro_::NERD1-GUS reporter (blue signal) is detected in inflorescence (A-B), the flower in stage 9 (C), the pistil in stage 8 (D), the pistil in stage 9 (E), mature pistil (F), mature flower (G), mature anther (H), and SAM and RAM of seedlings (I). Bars=25 μm.

### Overexpression of NERD1 increases plant productivity

We examined *NERD1* transcript levels in the low ovule number accession Altai-5 compared to Col-0. *NERD1* transcript accumulation was reduced in Altai-5 buds as compared to Col-0 but similar in Altai-5 and Col-0 leaves (Fig 7). We hypothesized that the low ovule number in Altai-5 may be linked to reduced *NERD1* expression in developing flowers. In order to determine whether increasing *NERD1* expression is sufficient to increase ovule number, we transformed Col-0 and the low ovule number accession Altai-5, with a *NERD1-GFP* fusion construct driven by the constitutively expressed Cauliflower Mosaic Virus 35S promoter (*35S::NERD1-GFP*). Overexpression of *NERD1* had no effect on ovule number in the Col-0 background, but significantly increased ovule number in the Altai-5 background (Fig 8A-C), indicating that the *NERD1* effect on ovule number is background-dependent. The *35S::NERD1-GFP* plants displayed an even more striking phenotype when overall plant architecture was examined (Fig 8A). In both the Altai-5 and Col-0 backgrounds, *NERD1* overexpression led to increased branching (Fig 8D) and shortened internode lengths between flowers, leading to an overall increase in flower number in the overexpression plants compared to untransformed controls (Fig 8E-H). Thus, *NERD1* overexpression leads to increased biomass and reproductive capacity, with up to a 2.5-fold increase in total flower number over the lifespan of the plant. While all independent transformants displayed increased branching and flower number, some of the *35S::NERD1* plants were male sterile (Fig S7). This male sterility correlated with *NERD1* expression levels and plants with higher *NERD1* transcript levels had more severe male sterility (Fig S7). The sterility effect was more severe in Col-0 than in Altai-5 (Fig S7). The lower endogenous *NERD1* expression in Altai-5 inflorescences might explain the lower sensitivity of Altai-5 to *NERD1* overexpression with respect to male fertility, demonstrating background-dependent sensitivity to *NERD1* levels for both ovule number and male sterility.

**Fig 7.**
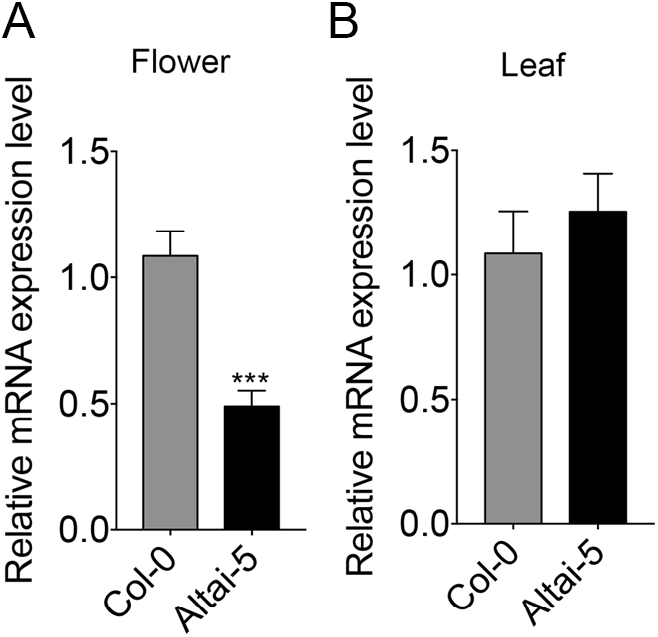
NERD1 expression is reduced in developing flowers of a low ovule number accession. (A) qRT-PCR of NERD1 in Col-0 and Altai-5 inflorescences. (B) qRT-PCR of NERD1 in Col-0 and Altai-5 leaves. “***” indicates statistical significance at *p* value<0.001 determined by Student’s t-test.

**Fig 8.**
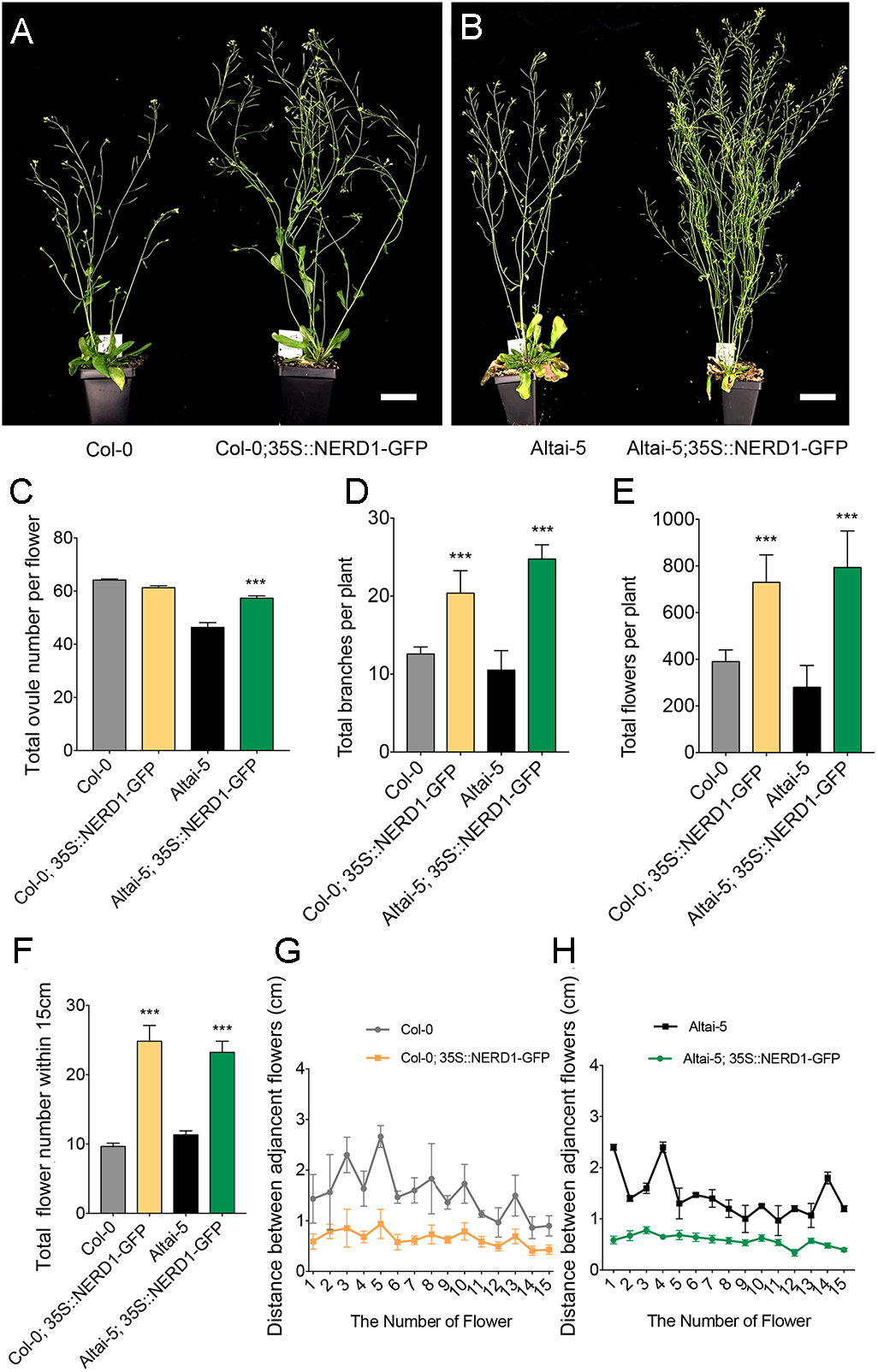
Overexpression of NERD1 affects plant architecture and fertility. (A-B) 35S::NERD1-GFP expressed in Col-0 and Altai-5 backgrounds. Bars=5cm. Bar graphs of total ovule number per flower (C), total branch number (D), total flower number per plant (E), number of flowers within 15 cm of the main shoot (F), and distance between adjacent flowers (G-H) in 35S::NERD1-GFP transformants in the Col-0 and Altai-5 backgrounds compared to untransformed controls.

*NERD1* was recently identified in enhancer screen performed on exocyst mutants with weakly dwarfed roots. *nerd1* mutants have reduced root growth as a result of impaired cell expansion, indicating that NERD1 may be a positive regulator of exocyst-dependent root growth(25). Consistently, the *35S::NERD1-GFP* plants had longer roots compared to the Col-0 control, indicating that root development is also sensitive to *NERD1* expression levels (Fig S8).

Three SNPs in NERD1 showed significant correlation with ovule number in our GWAS (Fig 9A-B). Two of them are synonymous SNPs that are not predicted to change the amino acid sequence, but the third is a non-synonymous SNP (C to A change in comparison to Col-0 reference) causing a Serine to Tyrosine change at amino acid 230 of NERD1 (Fig 9A-B). This non-synonymous SNP was present in 10 out of the 16 lowest ovule number accessions and not present in the 16 highest ovule number accessions (Fig 9A). Across all of the accessions in our GWAS panel, the “A” allele at this position was significantly associated with lower ovule numbers (Fig 9C). However, some accessions with the “C” allele of *NERD1* have low ovule numbers, including the Altai-5 accession (Fig 9A), underscoring that multiple mechanisms influence ovule number in Arabidopsis.

**Fig 9.**
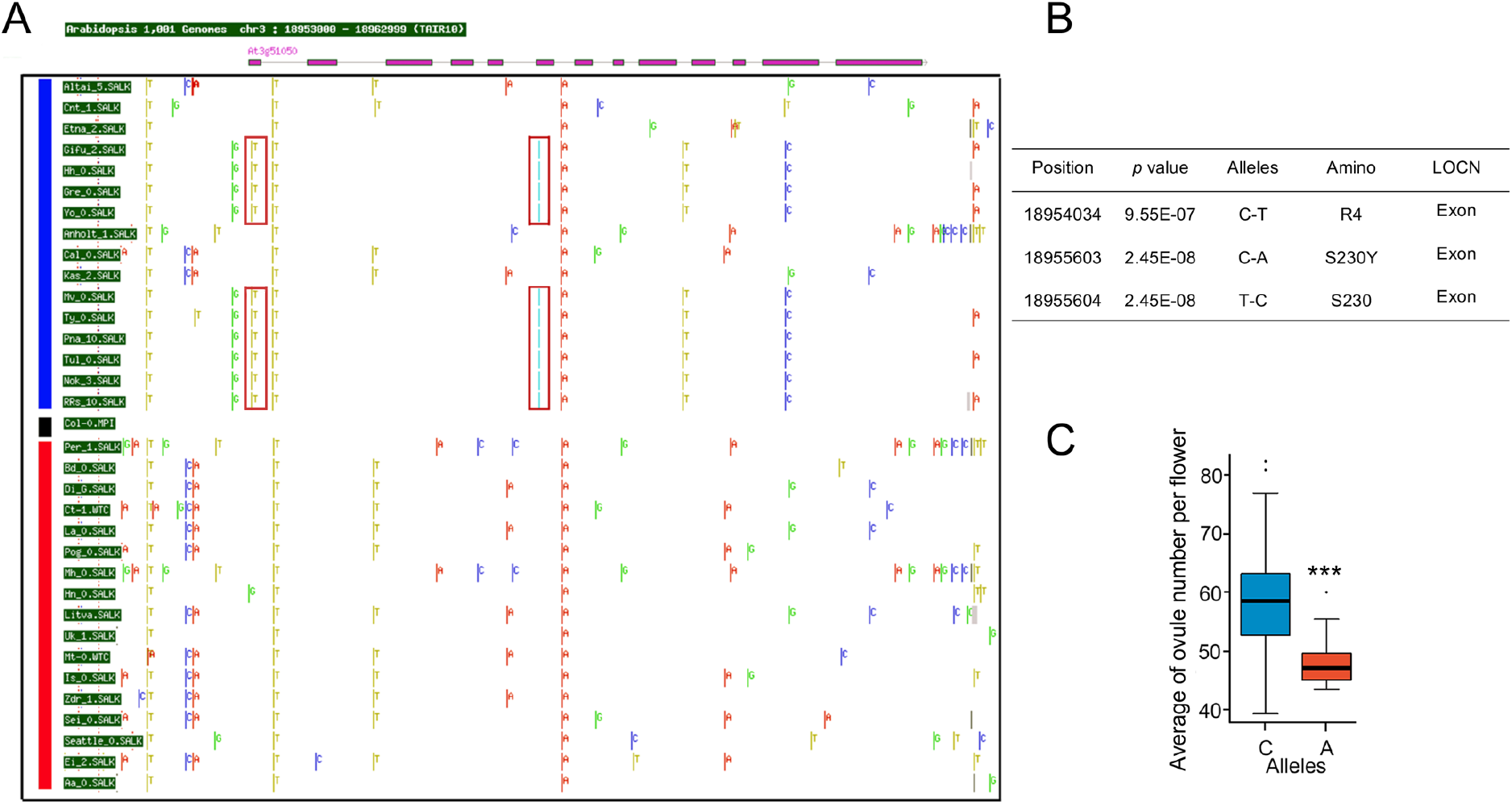
Sequence variation in the *NERD1* locus correlates with ovule number. (A) SNPs in and around the *NERD1* gene. The blue line indicates the 16 lowest ovule number accessions and the red line indicates the 16 highest ovule number accessions. SNPs are identified based on comparison to the Col-0 reference genome. The red boxes highlight low-ovule number associated SNPs (the turquoise bar indicates two SNPs in adjacent nucleotides, a non-synonymous C-A SNP and a synonymous C-G SNP). Only genomes that were available on the SALK 1,001 genomes browser (http://signal.salk.edu/atg1001/3.0/gebrowser.php) were considered. (B) One of the ovule number-associated SNPs in NERD1 causes a serine-tyrosine substitution in low ovule number accessions. (C) Average ovule number in *NERD1* “A” and “C”allele-containing accessions. The “A” allele is significantly associated with lower ovule numbers. “***” indicates statistical significance at *p* value<0.001 determined by Student’s t-test.

## Discussion

Our experiments revealed that Arabidopsis displays a wide variation in the number of ovules per flower, a key component of plant fitness and crop yield. Association genetics linked this natural variation to the plant-specific gene, *NERD1*, that regulates the number of ovules produced in Arabidopsis flowers and gametophyte development (Fig 3) and was previously found to alter root growth(25). Overexpression of *NERD1* led to dramatic effects on plant architecture, indicating that *NERD1* may be involved in regulating meristem activity during Arabidopsis development.

### GWAS reveals new ovule number-associated loci in Arabidopsis

A quantitative trait locus (QTL) mapping study utilizing variation between Ler and Cvi identified QTLs residing on chromosomes 1, 2 (near the *ERECTA* gene), and two QTL on chromosome 5(23). No follow-up study has identified the genes underlying these QTLs. Our population displayed a much larger range of ovule numbers per flower (39–84) than that seen in Cvi vs Ler (66 and 56, respectively). We identified 9 significant associations in our GWAS, but only one potentially overlaps with a known ovule number QTL (see chromosome 5 in Fig S2). Of the genes with an identified effect on ovule number, only BIN2 overlapped with a GWAS peak, indicating that our study has revealed at least seven novel regulators of ovule number in Arabidopsis. We molecularly identified two new genes that control ovule number that were linked to two GWAS peaks on chromosome 3. This expands that number of known ovule number determinants and confirms the value of GWAS as a forward genetics tool.

### NERD1 may transmit signals across membranes

*NERD1* is a plant-specific, low copy number gene that is found throughout the plant kingdom. NERD1 was recently identified as an enhancer of root development phenotypes in weak mutants of the exocyst subunits SEC8 and EXO70A1(25) While a direct interaction with the exocyst could not be identified, the authors identified Golgi localization of NERD1 and hypothesized that NERD1 could be involved in synthesizing or modifying pectins or other polysaccharide components of the cell wall that are synthesized in the Golgi and transported to the cell wall (25). Changes in cell wall elasticity mediated by pectin modifications have been correlated with lateral organ initiation at the shoot apical meristem (26, 27). The decreased ovule number in *nerd1* mutants and increased lateral branching and flower initiation in *35S::NERD1* transformants could be consistent with alterations in Golgi-synthesized cell wall components affecting cell wall mechanics and organ initiation.

Our GFP localization experiments confirmed Golgi localization of the majority of NERD1-GFP, but also identified fluorescence in the plasma membrane. In Arabidopsis, the NERD1 protein is predicted to encode a 698 amino acid protein with a signal peptide, one transmembrane domain, and a short (17 amino acid) cytoplasmic tail at the C-terminus. This topology is consistent with NERD1 being involved in transducing signals from the outside of the cell to the inside, perhaps as part of a complex with other membrane proteins. In animals, α-integrins and β-integrins share a similar topology with NERD1 and interact with each other to transduce information from the extracellular matrix to the inside of the cell(28). The existence of integrin-like complexes in plants remains controversial, since plants lack proteins with clear sequence homology to animal integrins. However, antibodies to animal β-integrins interact with plant proteins, indicating structural conservation(29). NERD1 has a predicted beta-propeller with a calcium-binding pocket similar to α-integrins in its extracellular domain(25), but determination of whether NERD1 functions in an integrin-like fashion awaits further investigation.

NERD1 could also interact with other types of proteins to transduce signals. In plants, receptor-like proteins (RLPs) with short cytoplasmic domains have been shown to heterodimerize with receptor-like kinases (RLKs) to form a receptor complex that transduces extracellular signals to intracellular signaling networks. The Arabidopsis genome contains 57 RLPs with Leucine Rich Repeats (LRRs) in their extracellular domains(30). These LRR-RLPs have a transmembrane domain near the C-terminus and a short cytoplasmic domain and are predicted to form complexes with LRR-RLKs. LRR-RLP/LRR-RLK complexes have been implicated in plant growth and development and plant immunity (reviewed in(31)). For example, the CLAVATA2 (CLV2) RLP forms a complex with the RLKs CORYNE (CRN) and/or CLAVATA1 (CLV1) RLK to perceive the CLAVATA3 (CLV3) ligand and trigger intracellular signaling related to shoot apical meristem maintenance(32–34). Likewise, the TOO MANY MOUTHS (TMM) RLP interacts with the LRR-RLKs ERECTA (ER) and ERECTA-LIKE1 (ERL) to regulate stomatal patterning(35, 36). In contrast to the LRR-RLPs, NERD1 does not share sequence homology with other proteins in the Arabidopsis genome. Distinguishing whether NERD1 functions similarly to RLPs, integrins, or some other signaling network will require identification of NERD1-interacting proteins in future work.

### *NERD1* and fertility

Homozygous *nerd1* mutants are completely male sterile are partially female sterile. This sterility is due to a lack of pollen production and problems in female gametophyte development leading to aborted embryo sacs. Transmission efficiency tests using heterozygous loss-of-function mutants revealed that *nerd1* could be transmitted through the egg at near 100% efficiency, indicating a sporophytic effect on female gametophyte development. Ovule development mutants that have defective integument development such as *short integuments 1* (*sin1*), *bell 1* (*bel1*) and *ant* fail to produce functional female gametophytes, indicating that female gametophyte development is dependent on properly differentiated sporophytic cells in the ovule(37–41). Sporophytic development in ovules seem to be normal in *nerd1* mutants, yet embryo sacs abort. As a membrane protein, NERD1 could transmit some unknown signal between sporophytic and gametophytic cells during embryo sac development.

The male fertility defect in *nerd1* plants is more severe than the female defect. Homozygous *nerd1* anthers are empty, indicating that pollen development is not initiated or aborted very early. Transmission efficiency tests using pollen from heterozygous *nerd1* plants crossed to wild-type females revealed that *nerd1* also has gametophytic effects on pollen function. The sporophytic effects could be related to early stages of anther development. In particular, specification of the tapetum is critical for pollen development (reviewed in(42)). In our *35S::NERD1* experiment, transformants that accumulated the most *NERD1* transcripts were male sterile. Together these results indicate that pollen development is sensitive to *NERD1* levels, i.e. either too much or too little *NERD1* is detrimental to pollen development. Future experiments should focus on determining the stage of anther and/or pollen development that is affected in *nerd1* mutants and the specific cell types that express *NERD1* in developing anthers.

Even though *NERD1* is expressed broadly throughout the plant, above ground vegetative development appears to be normal in *nerd1* loss-of-function mutants. However, *nerd1* roots are shorter than normal and have root hair defects that include bulging and rupture(25). *NERD1* could have distinct or related developmental functions in roots and flowers, as is seen for many of the genes involved in hormonal regulation of development(43).

### *NERD1* regulates lateral organ formation

*NERD1* overexpression under control of the constitutive 35S promoter dramatically changed plant architecture in both the Col-0 and Altai-5 backgrounds. *NERD1* overexpression plants produced significantly more branches than wild-type controls and produced significantly more flowers that were produced at closer intervals along the stem. Like branches and flowers, ovules are lateral organs produced from meristematic cells, indicating that *NERD1* may be a positive regulator of meristem activity. Lateral meristem activation and subsequent branching have been shown to be affected by several different hormones (reviewed in(44)). Classic experiments in *Vicia faba* showed that auxin produced in the shoot apex inhibits axillary meristems(45), and more recently strigolactones were identified as graft-transmissible suppressors of branching (reviewed in (46)). Gibberellins also repress shoot branching in Arabidopsis, maize, and rice, with GA-deficient mutants displaying increased branching compared to wild-type controls(47–50). In contrast, cytokinins and brassinosteroids are both positive regulators of branching(49–53). The mechanism through which these plant hormones work together to regulate branching is unknown, but auxin has been proposed to control cytokinin and strigolactone biosynthesis(54, 55), while the brassinosteroid signaling regulator *BES1* may inhibit strigolactone signaling to promote branching(56). The promotion of branching in *35S::NERD1* plants could be related to regulation of one or more of these hormonal pathways. Like *NERD1*, upregulation of both cytokinin and brassinosteroid signaling pathways have been shown to positively regulate branching and ovule number(15, 53, 56, 57), suggesting that *NERD1* may be intimately connected to these pathways. Future research is needed to explore the intersection between *NERD1* and hormonal pathways.

### *NERD1*-induced increases in ovule number are background-dependent

Overexpression of *NERD1* in the Col-0 and Altai-5 backgrounds led to increased branching and flower number, but ovule number was only increased in the Altai-5 background, suggesting a background-dependence on the ovule number trait. In Arabidopsis, natural accessions were shown to respond differently in their ability to buffer GA perturbations caused by overexpressing GA20 oxidase 1, which encodes a rate-limiting enzyme for GA biosynthesis(58). Genetic background dependence has been shown to be a wide-spread phenomenon in *C. elegans*, with approximately 20% of RNAi-induced mutations (out of 1400 genes tested) displaying different phenotypic severity in two different genetic backgrounds(59). Similar to our results with *NERD1*, where Altai-5 has lower endogenous levels of the *NERD1* transcript in developing flowers compared to Col-0, the severity of the *C. elegans* phenotypes could be linked to gene expression levels of either the target gene itself or other genes in the same pathway(59).

Our GWAS revealed multiple novel loci that control ovule number in Arabidopsis. It remains to be seen if exocyst function is linked to NERD1’s role in ovule number determination. Identification of NERD1 interacting proteins will provide candidates for other players in the NERD1 pathway. Some of the other novel loci identified in our GWAS may well participate in the same signaling pathway as *NERD1*. While *NERD1* is a promising candidate for engineering plants with increased ovule number, the differential responses to *NERD1* overexpression seen in Altai-5 and Col-0 suggest that specific alleles at other loci may be necessary for achieving maximum effect of *NERD1* overexpression.

## Materials and Methods

### Plant Material and Growth Conditions

*Arabidopsis thaliana* accessions and insertion mutants were ordered from the Arabidopsis Biological Resource Center at Ohio State University (ABRC). ABRC stock numbers for the accessions and insertion mutants used in our study are listed in supplementary tables 1 and 2. Seeds were sterilized and plated on ½ Murashige and Skoog (MS) plates. All plates were sealed and stratified at 4 °C for two days, and then transferred to the growth chamber (long day conditions, 16h of light and 8 h of dark at 22°C) for germination and growth. After one-week, seedlings were transplanted to soil. Many of the Arabidopsis accessions require vernalization for flowering(60), we therefore chose to vernalize all of the accessions in our study for 4 weeks at 4 °C. After vernalization, the plants were returned to the growth chamber and grown under the long day conditions described above. Seeds from transformed lines were sterilized and plated on ½ MS plate with 20mg/L hygromycin for selection of transgenic seedlings, which were then transplanted to soil and grown in long days.

### Ovule Number Phenotyping

For determination of ovule number per flower, we counted four to five siliques per plant and five plants per accession. Carpel walls were removed with a dissecting needle and total ovule number (including unfertilized and aborted ovules) was counted with the aid of a Leica dissecting microscope. To minimize age-related variation in ovule number, we counted siliques from flowers 6-10 on the primary shoot for all accessions.

### Genome-Wide Association Study

GWAS was performed using GWAPP, which is a GWAS web application for Genome-Wide Association Mapping in *Arabidopsis* (http://gwas.gmi.oeaw.ac.at) (22). In our study, 148 accessions had single nucleotide polymorphisms (SNPs) data available on the 1001 Full-sequence dataset; TAIR 9. Logarithmic transformation was applied to make the results more reliable for parametric tests. A simple linear regression (LM) was used to generate the Manhattan plot by using GWAPP(22). SNPs with P values ≤ 1 × 10^-6^ were further considered as candidate loci linked to alleles that regulate ovule number (a horizontal dashed line in Fig. 2 shows the 5% FDR threshold -log10p value=6.2, which was computed by the Benjamini-Hochberg-Yekutieli method). SNPs with <15 minor allele count (MAC) were not considered to help control false positive rates. 10 genes flanking the highest SNP for each locus were tabulated as candidate genes for each significant association.

### Cloning and Generation of Transgenic Lines

For complementation and overexpression experiments, Gateway Technology was used to make all the constructs. Genomic DNA fragments corresponding to the coding regions of candidate genes were amplified from either beginning of the promoter (defined by the end of the upstream gene) or the start codon to the end of the CDS (without stop codon) by PCR with primers that had attB1 and attB2 sites from Col-0 genomic DNA (see supplementary table 3 for primer sequences). For amplifying At3g60660 and At3g51050, PHUSION High-Fidelity Polymerase (NEB, M0535S) was used. All the PCR products were put into entry vector pDONR207 by BP reactions and then were recombined into destination vector pMDC83 (GFP) by LR reaction(61).

Native promoter constructs were amplified from ~2KB promoter region to the end (without stop codon) by PCR with primers that had attB1 and attB2 sites from Col-0 genomic DNA. The PCR products were put into entry vector pDONR207 by BP reactions and then were recombined into destination vector pMDC32 (tdTomato) and pMDC163 (GUS) by LR reaction. All constructs were transformed into *Agrobacterium tumefaciens* strain GV3101 and then used for plant transformation by the floral-dip method(62).

### GUS Staining

GUS staining was performed as previously described(63). Samples were imaged with Differential Interference Contrast (DIC) on a Nikon Eclipse Ti2-E microscope. 12 independent T1 NERD-GUS transformants were analyzed and showed similar GUS patterns.

### Transient Expression in *N. benthamiana*

Leaves from 3-4 week old *N. benthamiana* were co-infiltrated with 35S::NERD1-GFP and Golgi-mCherry, ER-mCherry, PM-mCherry, Plastid-mCherry and Peroxisome-mCherry markers from(64) as previously described(65). Leaves were imaged 2-3 days after infiltration with a Nikon A1Rsi inverted confocal microscope under 20x dry and 40x water objectives with GFP excited by a 488nm laser and mCherry excited by a 561nm laser in normal mode.

### Alexander Staining of Pollen

Mature anthers from Col-0 and *nerd1-2/nerd1-2* were dissected under the Leica dissecting microscope and placed into a drop (20 μl) of Alexander staining solution on a microscope slide(66). After several minutes of staining, samples were imaged with a Nikon Eclipse Ti2-E microscope.

### Quantitative Real-time RT-PCR

For qRT-PCR, leaves and young flowers were collected from mature plants of Col-0 and Altai-5. Samples were immediately frozen in liquid nitrogen, ground, and total RNA was extracted using the E.Z.N.A Plant RNA kit (OMEGA, USA). Oligo-dT primers and Superscript II reverse transcriptase (Invitrogen) was used for cDNA synthesis. qRT-PCR reactions were prepared using SYBR Green PCR Master Mix and PCR was conducted with a StepOnePlus RT-PCR system. Relative quantifications were performed for all genes with the Actin11 used as an internal reference. The primers used for qRT-PCR shown in supplementary table 3.

### Phylogenetic Analysis

The cladogram tree was generated in MEGA7, which nucleotide distance and neighbor-join tree file were calculated by PHYlogeny Inference Package (PHYLIP, version 3.696). The phylogenetic tree of NERD1 was inferred using neighbor-joining method in MEGA7(67). The associated taxa clustered together with the bootstrap test (1000 replicates)(68). All the branch lengths are in the same units as those of the evolutionary distances used to generate the phylogenetic tree.

## Supplemental Material

**Supplementary Figure 1.** Statistical analysis of the association between geographic distribution and ovule number per flower in the Arabidopsis accessions used in the GWAS.

**Supplementary Figure 2.** Previously identified ovule number related genes’ position in relation to the Manhattan plot generated from the ovule number GWAS.

**Supplementary Figure 3.** Heat map of candidate gene expression patterns in reproductive tissues.

**Supplementary Figure 4.** Average ovule number per flower of candidate gene T-DNA mutants on chromosome 3.

**Supplementary Figure 5.** Homozygous *nerd1* mutants have high levels of infertility can be partially rescued by pollinating with Col-0 wild-type pollen.

**Supplementary Figure 6.** Co-localization of 35S::NERD1-GFP with peroxisome and plastid markers.

**Supplementary Figure 7.** Male sterility in *35S::NERD1* plants is linked to high *NERD1*expression.

**Supplementary Figure 8.** *NERD1* is involved in in root growth.

**Supplementary Figure 9.** Amino acid changes in and around the *NERD1* gene in low and high ovule number accessions.

**Supplementary table 1.** List of Arabidopsis accessions used in the study and their ovule number data.

**Supplementary Figure 2.** Chromosome 3 candidate genes and insertion mutants.

**Supplementary Figure 3.** List of primers used for genotyping and cloning.

## Acknowledgements

We thank Dr. Brian Dilkes, Dr. John Fowler, Dr. Daniel S. Jones, Thomas Davis, Yan Ju, and Rachel Flynn for helpful discussions and comments on the manuscript. We thank Dr. Xutong Wang for help with the statistical analysis. Purdue University Start-up funds and a grant from the Oklahoma Center for the Advancement of Science and Technology #PS14-008 to S.A.K. supported this work.

## Author contributions

**Conceptualization:** JY SAK.

**Data curation:** JY.

**Formal analysis:** JY SAK.

**Funding acquisition:** SAK.

**Investigation:** JY.

**Methodology:** JY.

**Project administration:** SAK.

**Resources:** JY SAK.

**Supervision:** SAK.

**Validation:** JY.

**Visualization:** JY.

**Writing - original draft:** JY.

**Writing - review & editing:** JY SAK.

